# Panaconda: Application of pan-synteny graph models to genome content analysis

**DOI:** 10.1101/215988

**Authors:** Andrew S. Warren, James J. Davis, Alice R. Wattam, Dustin Machi, João C. Setubal, Lenwood S. Heath

## Abstract

**Motivation:** Whole-genome alignment and pan-genome analysis are useful tools in understanding the similarities and differences of many genomes in an evolutionary context. Here we introduce the concept of pan-synteny graphs, an analysis method that combines elements of both to represent conservation and change of multiple prokaryotic genomes at an architectural level. Pan-synteny graphs represent a reference free approach for the comparison of many genomes and allows for the identification of synteny, insertion, deletion, replacement, inversion, recombination, missed assembly joins, evolutionary hotspots, and reference based scaffolding.

**Results:** We present an algorithm for creating whole genome multiple sequence comparisons and a model for representing the similarities and differences among sequences as a graph of syntenic gene families. As part of the pan-synteny graph creation, we first create a de Bruijn graph. Instead of the alphabet of nucleotides commonly used in genome assembly, we use an alphabet of gene families. This de Bruijn graph is then processed to create the pan-synteny graph. Our approach is novel in that it explicitly controls how regions from the same sequence and genome are aligned and generates a graph in which all sequences are fully represented as paths. This method harnesses previous computation involved in protein family calculation to speed up the creation of whole genome alignment for many genomes. We provide the software suite Panaconda, for the calculation of pan-synteny graphs given annotation input, and an implementation of methods for their layout and visualization.

**Availability:** Panaconda is available at https://github.com/aswarren/pangenome_graphs and datasets used in examples are available at https://github.com/aswarren/pangenome_examples

**Contact:** Andrew Warren

## 1 Introduction

Research in comparative microbial genomics has largely been organized around the concept of reference genomes (Tatusova *et al.*, 2014). Reference genomes provide a useful comparative touchstone for closely related organisms. However, they may not represent the biological diversity of an entire group of genomes. Currently there are more than 96,000 publicly available bacterial genomes sequenced, and this number is rapidly increasing (Wattam *et al*., 2016). Some closely related species have large numbers of sequenced genomes, creating interesting comparative challenges e.g. *E.coli* has more than 5,400 isolates while *S. aureus* has almost 9,000. As this sampling through sequencing becomes both deeper and broader, reference genome based methods become less effective at characterizing groups of organisms.

Pan-genomes represent a wide range of methods to analyze groups of genomes as a whole. One basic representation is the unique and core gene sets of a group (Vernikos *et al*., 2015, Page *et al*., 2015). This method provides a general characterization of content but does not capture the inter-relations between members of a group nor does it model the genomic context in which the gene sets occur. Other more detailed methods create a pan-genome representation at the nucleotide level. One such method, SplitMEM (Marcus *et al.,* 2014), creates a compressed de Bruijn graph for representing shared variation and single nucleotide polymorphisms. Strategies are still evolving to fully leverage this complex representation. The pan-synteny model targets genome architecture analysis and thus can be placed somewhere on the meso-scale of pan-genome analysis methods, being more detailed than a simple categorization of gene types and more abstracted than a SNP level analysis.

Whole genome alignment methods also take on the challenge of producing a detailed genome comparison at the nucleotide level. These methods can assist greatly when performing SNP and phylogenetic analysis. Many methods require the specification of a reference genome, relatively high memory, and that the genomes be closely related. One example of this, Harvest, is a software package for establishing core genome alignments between multiple genomes that recruits highly related genomes using the genome distance metric MUMi (Deloger *et al*., 2009). There are a number of methods available for performing whole genome alignment (Earl *et al*., 2014), many of which are useful for the creation of SNP trees and analyses for comparing closely related genomes at the DNA level. Lee *et al*., 2002 introduces the Partial Order Alignment (POA) graph for representing alignments and calls into question the practice of representing multiple sequence alignments as grids, since they are not well adapted for representing complex rearrangements and gaps accurately. Raphael *et al*., 2004 follows this up by introducing the ABA graph constructed to align proteins given initial BLAST seed alignments. The method detailed here also performs a whole genome multiple sequence alignment at the gene order level and encodes it as a pan-synteny graph.

Comparative methods that focus on synteny traditionally focus on finding syntenic blocks. As Ghiurcuta and Moret, 2014 point out, homology forms the basis for the construction of most synteny block analysis and the creation of blocks usually involves the process of “extending homologies among markers to homologies among blocks.” A syntenic block is typically thought of as conserved ordering of homologous genes between two or more genomes. DRIMM-Synteny (Pham and Pevzner, 2010) is a method for finding synteny blocks in multiple mammals using alignment de Bruijn graphs (A-Bruijn graph). An A-Bruijn graph is a generalization of a de Bruijn graph that represents alignments through coincident labelling of sequence position on edges and nodes. DRIMM uses the creation of an A-Bruijn graph and its subsequent processing to approximate the minimum number of edits necessary to convert from one sequence to another in the pursuit of synteny block construction. This is done by constructing or finding non-branching paths in an A-Bruijn graph with multiplicity greater than one.

Sibelia (Minkin *et al*., 2013) is a follow-on project to DRIMM and also uses A-Bruijn graphs to construct synteny blocks. Sibelia uses an iterative refinement approach based on different *k*-mer sizes in order to represent synteny blocks in a hierarchical structure at different levels of granularity. Both the Sibelia and DRIMM approaches edit sequences to remove loops (also known as whirls) and bulges in a de Bruijn graph to achieve non-overlapping synteny decompositions. In doing so, it destroys sequence continuity and limits the discovery of potentially complex evolutionary relationships. The pan-synteny graph approach we outline here bypasses the need to remove these anomalies and preserves sequence continuity.

## 2 Algorithm

While many high level pan-genome analysis methods use protein families to describe group content, there is no restriction by Panaconda to protein coding genes or protein families. This form of analysis only requires that the units in question be given as contiguous intervals situated on a sequence and that those units have a sequence and genome designation and that they be shared across genomes, e.g. a protein family or an orthologous group. We refer to the units to be analyzed as features and to their groupings as families. For a given Panaconda run, let *G* be the set of all genomes in an analysis, *P* be the set of all sequences in those genomes, and *F* be the set of all features. Pan-synteny graphs are an undirected graph model of both gene neighborhood and synteny for multiple organisms. For the set of sequences *P* included in a pan-synteny graph *R*, each sequence *p* ∈ *P* of length *n* is represented as a path in the pan-synteny graph, where each feature with a family assignment is represented by a node along that path. More formally, if a genome sequence *p* ∈ *P* is represented by a sequence of features *f*_1_, *f*_2_,…, *f*_*n*_, and these features label a set of nodes *v*_1_, *v*_2_,…,*v*_n_ in the pan-syteny graph such that there is a bijection from features to nodes representing *p*, then every pair of consecutive labels *f*_*i*_ and *f*_*i*+1_ will label adjacent vertices in the pan-synteny graph such that a path is constructed. We refer to the expression of each sequence in *P* as a path in *R* as the full-path property.

Similar to DRIMM and Sibelia the Panaconda method creates a de Bruijn graph as part of the conserved gene order analysis. Unlike these two approaches, Panaconda processes the de Bruijn graph so that sequence continuity is preserved in the resulting simplified pan-synteny graph (PS-graph). Panaconda takes as input an annotation, ordered by lower base coordinate, of genome features with some family designation. These annotations are processed into k-mers of family assignments and are used to construct a labeled de Bruijn graph that we designate a *reverse fragment graph* (RF-graph). An RF-graph is a form of double stranded de Bruijn graph (Lin *et al*., 2014) *DB*\*(*G, k*) that encodes both the forward and reverse strand of a sequence on the same node and maintains four edge types to model the transition from one *k*-mer orientation to another.

Panaconda takes a number of parameters that govern how the PS-graph is constructed. Let *m* be the minimum multiplicity of sequences that must be incident on a node or edge for it to be shown in the PS-graph. Let *k* be the *k*-mer size given to the de Bruijn graph construction that breaks each sequence of size *n* into *n* − *k* + 1 fragments. The *k*-size can be viewed as the minimum size of conserved neighborhood to constitute a syntenic block.

### 2.1 RF-graph creation

The RF-graph and its subsequent processing leverages the edge types found in a double stranded de Bruijn graph and maintains a multigraph model, similar to the A-Bruijn graph found in Sibelia, such that the features *f*_1_ *f*_2_,…*f*_*n*_ in a given *p* ∈ *P* label a set of directed edges in the RF-graph and that those edges can have more than one label from one or more sequence. As in Pevzner *et al*., 2004, we refer to the number of features that label a given node as its multiplicity. As part of the rf-node creation process a hash function is applied to k-mers, which performs a lexicographic comparison of the beginning and ending characters of the k-mer. If the character at the end is less than the beginning, the k-mer is reversed. If the beginning and end character are equivalent then the adjacent characters are compared using the same criteria. This continues inward until an inequality is found that determines the orientation or the middle of the *k*-mer is reached in which case the orientation remains unchanged from its source genome. This enforces a consistent ordering of k-mer and its reverse, and insures that a k-mer and its reverse are hashed to the same rf-node. This enables genomes to be placed in correct syntenic context within the RF-graph regardless of the direction of sequencing and subsequent ordering of feature families. Because we want to maintain sequence continuity in processing the RF-graph and the downstream PS-graph, each of the *n*−*k*+1 transitions between k-mers of a given sequence *p* ∈ *P* of length n is modeled as a distinct set of *n* − *k* +1 edges or edge labels. This means that for a given sequence the transition from one position to the next can be specifically referenced when traversing the graph.

Each bin in the resulting hash instantiates an rf-node. Edges in the RF-graph are created between RF-nodes that represent neighboring k-mers in the sequence from a source genome (Figure 1). The RF-edges are directional, taking on the direction from the source k-mer to the next one in sequence. Each rf-node is annotated with the features *f* ∈ *F* from the genome(s) of origin and nucleotide coordinates of those features such that the original sequence of features or feature families can be recovered for any given *p* ∈ *P*.

**Fig. 1.**
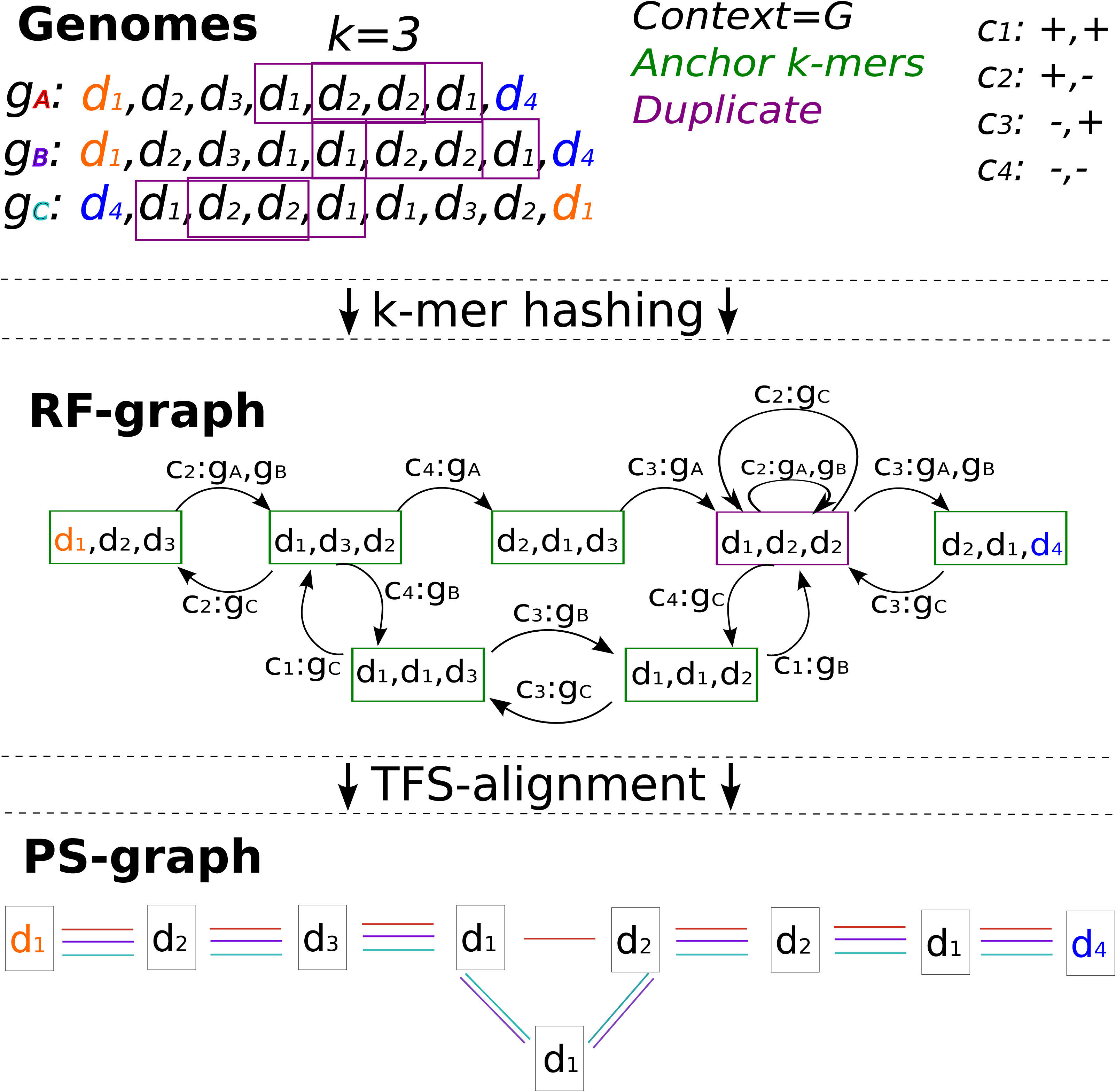
The RF-graph and resulting PS-graph given three sequences, *k* = 3, and *C_G_*. Although not shown explicitly here, the RF-edges can be traversed in reverse. In this case the sequence is effectively reversed and the complement of the edge type is used.

To account for the reversal of some k-mers, RF-edges are assigned categories to distinguish the orientation of its incident k-mers in the original sequence. A *k*-mer’s reversal relative to the original genome sequence is represented with a – and the absence of reversal with +. The RF-edge categories can then be given as directed edges in the form {(+, +), (+, −), (−, +), (−, −)}. Due to the constraints of the reversing hash function and directional edges, only 1 category can exist per directed edge. When traversing from one RF-node to another, RF-edges can be used to determine the orientation of the next k-mer and which side of the *k* − 1 ovelap will be appended in the next RF-node. The labels and their categories are used in subsequent transformation from an RF-graph to a PS-graph.

After an RF-graph is formed, it is processed by an algorithm we call Thread First Search alignment (TFS-alignment). The TFS-alignment procedure creates a whole genome alignment, at the granularity of feature families, as a labelled, undirected PS-graph. The PS-graph, is an expansion of the *k*-mers, represented by RF-nodes, into their *k* constituent components, and is labelled by the TFS-alignment procedure to represent the position of individual features from the original sequences with their grouping forming the basis of alignment (see Figure 1). Previous comparative genomic analysis of synteny has focused on determining synteny between genomes, between contigs in the same genome, within contigs, or some combination thereof [32, 10, 5, 3, 27]. In pan-synteny graph construction, the syntennic context, *C*, defines the relationships of interest that should underpin a synteny block. If the context is at the genome level, *C*_*G*_, then a synteny block is defined by similarity between genomes and not within them. If the context is between-contigs, *C*_*P*_, a synteny block is defined by similarity between genomes, between contigs, and not within contigs. Finally, if the target context is within-contigs, represented as *C*_Ø_, then synteny blocks represent similarity within contigs, between contigs, and between genomes.

The context given as an input parameter to Panaconda governs which nodes in the RF-graph are used as anchors to begin an alignment. In order to create synteny blocks in accordance with the syntenic context, pan-synteny graph construction uses context bins. For a given context {*C*_*G*_, *C*_*P*_, *C*_Ø_} a context bin *c* is a grouping of sequences from *P* used to determine if a *k*-mer and its corresponding rf-node are marked as *duplicates*. If the context is set to the genome level, *C*_*G*_, then each *g* ∈ *G* is a context bin. If the context is set to the sequence level, *C*_*P*_, then each *p* ∈ *P* is a context bin. If the context-level is within-contigs, *C*_Ø_, then there is no context bin. If a *k*-mer occurs more than once within any context bin then its corresponding rf-node is marked as a *duplicate*. If a k-mer is a palindrome then its corresponding rf-node is also labeled as such. All rf-nodes that are neither duplicates nor palindromes are *anchor* nodes (see Figure 1). *Anchor* nodes serve as the “gluing” agent to begin an alignment in the TFS-alignment procedure which we present here.

### 2.2 PS-Graph Creation

#### 2.2.1 TFS-Traversal

Starting from an anchor node in the RF-graph, we employ a search strategy, which we refer to as thread first search (TFS). The thread first search algorithm assumes a multigraph *M*(*V, E*) view of an RF-graph labeled with sequences from a set *P* where a given *p* ∈ *P* of length *n* corresponds to *n* labels *f*_1_, *f*_2_,…, *f*_*n*_ such that each is assigned to a node in *M*. In this case every node in *M* is labeled with at least one label from a *p* ∈ *P*. For a sequence of features *f*_1_, *f*_2_,…, *f*_*n*_ let the pair *f*_*i*_ and *f*_*i*+1_ represent adjancent features in the sequence for 1 ≤ *i* ≤ *n*. Following the ordered labels of each element of a sequence *p* ∈ *P* corresponds to a walk of the graph. We refer to the set of edges that correspond to a walk of a given sequence as a thread. Anchor nodes are labeled according to the context bins previously defined.

In the TFS algorithm nodes are queued for visiting based on the feature labels and their corresponding threads. On visiting an anchor node, all threads with features present in that node are *activated.* If a thread in a node is *active* then it’s edges incident to the current node are used to queue adjacent nodes. For a given visit, nodes can only be queued once, though more than one active thread may induce its queueing. For a queued node *v*_*q*_ all threads that induce the queueing of *v*_*q*_ are grouped together and passed on to the visit of *v*_*q*_. We call this grouping of threads a *bundle.* Bundles are necessary because in TFS traversal, non-anchor nodes do not activate threads. Thus only threads that are pre-activated by way of a bundle are queued on a visit to a non-anchor node.

Starting at an anchor node, TFS *activates* all threads present in that node and adds to the visit-queue those adjacent nodes that have connecting edges from active threads. Because a visit to a non-anchor node only queues those adjacent nodes whose threads are already active in the incoming bundle, an anchor node only queues adjacent nodes once and a non-anchor node may be visited and queue adjacent node visits multiple times. The addition of a thread to a bundle via a feature *f_i_* is encoded in a two step process. First the next feature *f*_*i*+*k*−1_ or *f*_*i*−*k*−1_ that is to be added at one end of the *k* − 1 overlapping characters is projected according to one of the four RF-edge types. The projected feature is then added to a feature packet which is passed forward to the visit of *v*_*q*_.

For a node *v* and k-mer index *x*, where 1 ≤ *x* ≤ *k*, let *Fv*_*x*_ be all features at position *x* in the k-mer represented by *v*. For a feature *f*_*i*_ at position *i* of a sequence *p* let *V*^+1^(*f*_*i*_, *v*) give the adjacent feature 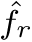, and corresponding node 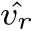, that shifts the reading frame for *v* in the sequence *p*, containing *f*, to the right. Likewise let *V*^−1^(*f*_*i*_, *v*) give the adjacent feature 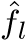 and corresponding node 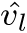 that shifts the reading frame to the left. For a given RF-graph, we give the following detailed procedure for thread first search.

1. **function** TFS(*M*, *v*, *v*_*p*_, *T*)
  - INPUT: *M* an RF-graph with anchor, duplicate, and palindrome labels
  - INPUT: Node currently being visted *v* ∈ *V* and thread bundle *T*
  - For a node *v* being visited let *v*_*P*_ be the previous node.
2. Queue = Ø
3. **if** *v* is an anchor node **then**
4. *F*′ = *F*(*v*_*k*_) ∪ *F*(*v*_1_)
5. **else**
6. *F*′ = *T*
7. **for** Features *f*_*i*_ ∈ *F*′) **do**
8. 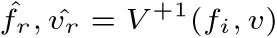
9. 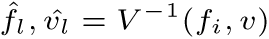
10. **if** 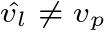 and *f*_*i*_ not exhausted in *v* **then**
11. **if** 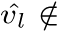 Queue **then**
12. *T*′ = {*f*_*l*_}
13. Append (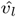 *T*′) to Queue
14. **else**
15. Update Queue(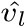, *T*′) by adding *f*_*l*_ to *T*′
16. **if** 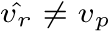 **then**
17. **if** 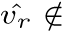 Queue **then**
18. *T*′ = {*f*_*r*_}
19. Append (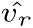 *T*′) to Queue
20. **else**
21. Update Queue(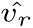 *T*′) by adding *f*_*r*_ to *T*′
22. Mark *f*_*i*_ as exhausted in *v*
23. **while** Queue 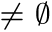 **do**
24. (*v*_*n*_, *T*′) = Pop Queue
25. TFS(*M*, *v*_*n*_, *v*, *T*′)

#### 2.2.2 TFS-Alignment

To create a pan synteny graph, Panaconda runs a process designated TFS-alignment. In the manner of the ABA-alignment algorithm (Pevzner *et al*., 2004) and the POA algorithm (Lee *et al.,* 2002), the TFS-alignment process “glues” features together by emitting a new node in the PS-graph with annotations that indicate the features being aligned in that node. Given an RF-graph for a set of genomes *G*, the TFS-alignment visits each node according to TFS-traversal and emits connected nodes in the PS-graph. Depending on the context *C* and the resulting anchor nodes, alignments will be started between sequences in different genomes (*C*_*G*_); between sequences in both the same genome and different genomes (*C*_*P*_); or between and within sequences in the same or different genomes (*C*_Ø_) (see Figure 2).

**Fig. 2.**
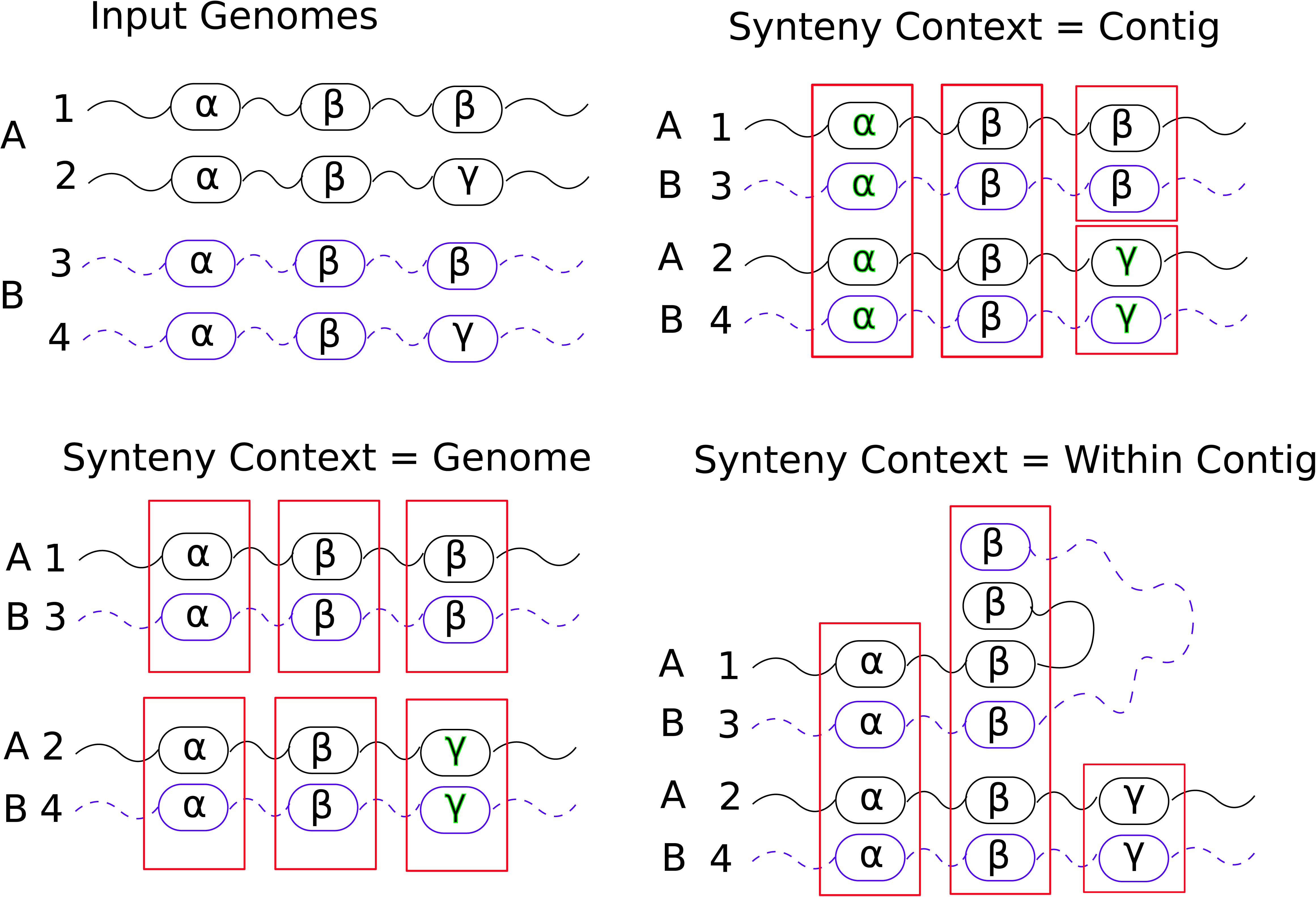
The synteny context is given as input to the TFS-Alignment algorithm and dictates how alignments are started. Here the red boxes show the default alignments in each case and correspond to the PS-nodes to be created.

TFS-alignment is performed such that, by default, all features underlying the same character position in an rf-node are assigned to the same PS-node. As we shall make clear subsequently, it is not always possible to do this and maintian the full-path property of the PS-graph. Another property of the PS-graph is that each feature *f* ∈ *F* given by *G* is only annotated to a single node in the PS-graph. The TFS-Alignment algorithm can be started on any available anchor node. In Panaconda, the TFS-alignment is started iteratively on anchor nodes in decreasing order of thread multiplicity. We refer to the process of visiting an rf-node and creating PS-nodes as *expansion.*

By assigning features to the same PS-node, we say they are *glued* together. In the default case, the first rf-node visited is expanded to *k* PS-nodes. For a given traversal, after the first node only the *k*th family of an adjacent rf-node outside the k − 1 overlap need be expanded unless new threads are activated. We refer to this position in the k-mer as the *target position*. Which side of a k-mer the target position occurs on can be determined by the RF-edge class traversed. The target position is always used to determine which nodes to queue next.

As detailed in the TFS-traversal procedure, a new thread is activated at an anchor node when it is incident on that node but not part of the incoming bundle. When a new thread is activated, all *k* features in that thread are assigned to PS-nodes, *k* − 1 of those to existing PS-nodes for the *k* − 1 pre-processed positions. In this case, both the target position and the position on the other side of the k-mer are used to queue nodes for subsequent visits. If the node to be queued represents the previous node visited, then the bundle information is returned in the unwinding phase of the TFS-traversal, similar to that of depth first search. Figure 1 gives an example of the TFS-alignment conversion from an RF-graph to a PS-graph. An RF-graph can be viewed as a specially labeled double stranded de Bruijn graph, *DB*\* (*G*, *k*) where, *k* ≥ 3, with *k* feature family labels. The TFS-alignment can then be viewed as a conversion, under certain constraints, to a graph with a single family labeling nodes and an edge representing a neighbor relation from a block of at least *k* characters (or nodes) with multiplicity of *m* or higher.

#### 2.2.3 Collisions

As part of the alignment process PS-nodes are created and annotated with the feature, sequence, and genome given as input to Panaconda. Due to complex patterns of conservation and shuffling, it sometimes happens that TFS-alignment, in accordance with the bundling information generated at anchor nodes, will attempt to assign the same genome or sequence in violation of the given context *C*_*G*_ or *C*_*S*_. In essence, the alignment instructions generated at two different anchor nodes conflict with each other under the constraints of the context *C*. We refer to this conflict in alignment information as a *collision.* We show here that collisions are a valuable source of information for the recognition of evolutionary relationships. Unless otherwise stated, it is safe to assume for the purposes of this explanation that the context is set to be “between genomes” *C*_*G*_.

In the default case, features sharing the same position in an anchor k-mer are assigned to the same PS-node. Due to short repeat sequences, it is not always possible to both group all features sharing a position in an anchor k-mer and maintain the full-path property of the PS-graph. We refer to this situation as the “shift” problem. The shift problem is caused by repeat characters and the solution is analogous to creating a gap in the alignment representation of a sequence to accomodate an extra repeat character in another.

As seen in Figure 3, the third character in sequence p1 (reading left to right) can be “shifted” to the right in the alignment if the rf-node corresponding to *α*3, *α*4, *α*5 is processed first. Shifts are often detected when activating a new thread at an anchor node. As a consequence of the TFS-traversal algorithm passing forward only bundle information that is relevant in the adjacent node, threads that have already been processed can be “rediscovered” when visiting an anchor node further in the traversal process. In this case, a rediscovered thread and its corresponding features may have already been partially assigned. The TFS-alignment algorithm treats the corresponding PS-nodes to which these features have been previously assigned as available positions for alignment for the features of all newly activated threads. Continuing with the example in Figure 3, if the path in the RF-graph that corresponds to PS-edge (*x*3, *x*6) is taken first, then the threads for *p*_2_ and *p_3_* are rediscovered and activated. When this happens, the TFS-alignment algorithm will investigate an alignment to a previous instance of the repeat character in the same sequence. In such cases the context constraint determines if this is viewed as a collision. In the case where the path corresponding to *p*_2_ is taken first, a shift will still occur relative to *p*_3_. In both these cases the TFS-alignment emits an extra node to represent the extra character and preserve the full-path property. If the highest repeat path is traversed first then three nodes are emitted with no special exception needed and only the direction of approach determines the resulting shift. We note that based on varying traversal order, the aligning and emitting of nodes may result in alignment graphs that are not isomorphic. Because threads are only activated at anchor nodes the default expansion step in TFS-alignment only creates one PS-node per position in the *k*-mer underlying an RF-node. However, due to the extra PS-node emission during the shift handling procedure it is possible for multiple PS-nodes to be available for feature assignment at a fixed position in an anchor k-mer. In this case all emmitted PS-nodes are assigned an identical “alternate ID” in the output expressing this group relationship. We leave the possibility of treating these groups as a kind of hypernode in the pursuit of isomorphic alignments as a subject for future development.

**Fig. 3.**
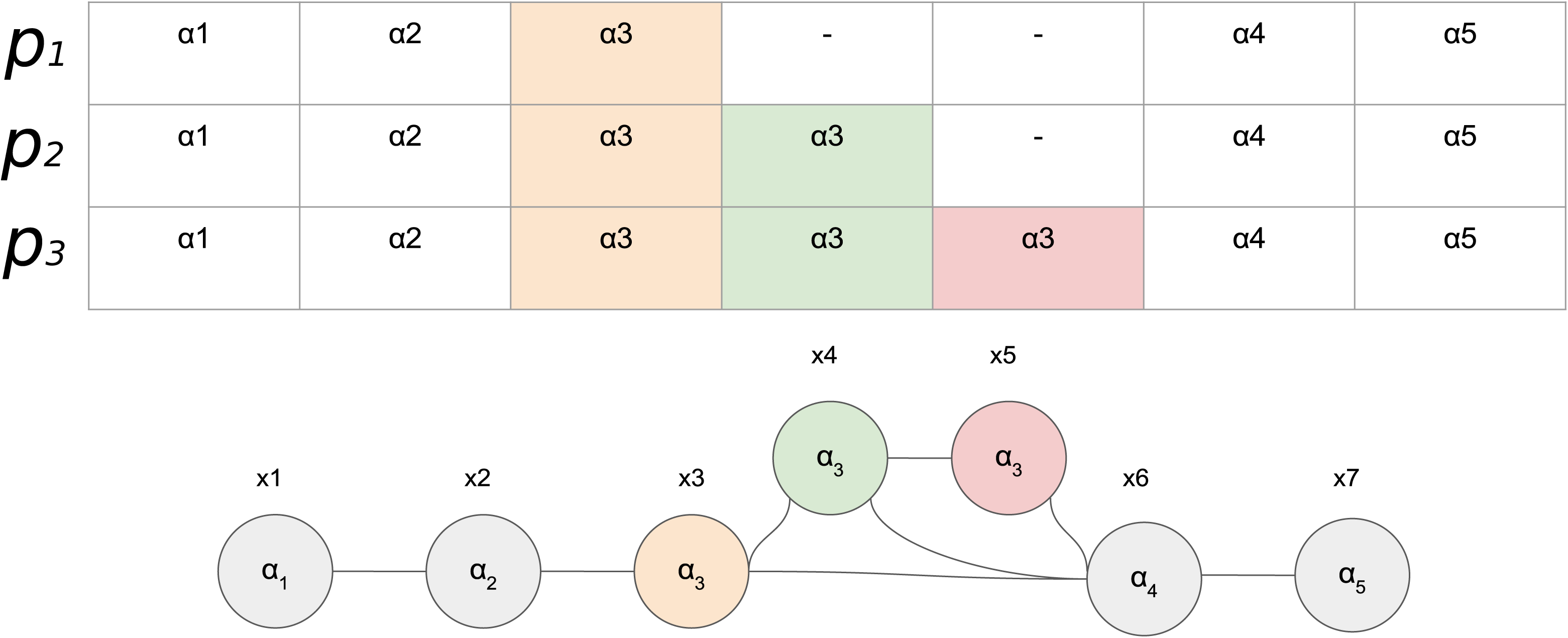
Due to repeat sequences alignments can be shifted giving variable annotation to resulting PS-nodes. The top shows the given sequences in a traditional multiple sequence alignment arrangement (from which the configuration of the RF-graph can be determined). The bottom is a possible PS-graph if *k* = 3.

The shift problem and resulting extra nodes turns out to have large downstream effects. As previously described, when new threads are activated a portion of the k-mer at the rf-node will represent pre-processed positions and will have corresponding PS-nodes at each position. Because repeats can “stack up” at a given location in the alignment a choice arises as to which PS-node to assign an activated feature to. In this case, the PS-nodes being considered all represent the same feature family. Panaconda is designed to construct alignments such that features from similar sequences are assigned together. For this purpose we introduce the concept of an *instance key.* For a given feature *f* an instance key *I*_*key*_ (*f*) has two components. The first component we refer to as the *i-component* and is a set representation of all rf-nodes (k-mers) in which a feature occurs. Depending on the position in its source sequence, a feature *f* may have an i-component from size 1 to *k.* Let all rf-nodes *V* in an RF-graph *M* (*V*, *E*) with *n* number of nodes | *V|* = *n* be assigned an integer ID value 1 to *n*. Let *f_t_* be a feature *f_t_* ∈ *F* from a sequence *p* ∈ *P* that occurs in rf-nodes with ID’s *h, i, j*; then the i-component of it’s instance key can be represented as {*h, i, j*}.

The second component of an instance key, the r-component, is the length of the character repeat, if any, in which the feature occurs in p. The r-component is encoded in such a way that its value is disjoint from the set of possible values for the i-component. Let the same feature *ft* occur in a repeat of size *y* in sequence p, and it’s r-component be represented as *Rt(y).* Given a subsequence *F*_*i*_, *f*_2_, *f*_3_, *f*_4_, *f*_5_, *f*_6_ of *p* ∈ *P* that corresponds to a character representation *d*^1^, *d*^2^, *d*^2^, *d*^2^, *d*^3^, *d*^4^, such that the *I*th position in one corresponds to the *I*th position in the other: we give the r-component value of feature *F*_2_ as *Rt*(*F*_2_) = 3_*repeat*_. An instance key is the union set of these two components such that for *ft*, *I*_*key*_ (*f*_*t*_) = {*h*,*i*,*j*,*Rt*(*y*)}.

For a visit to an rf-node *v* in which new threads are activated, let 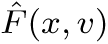(*x*, *v*) be those features at a fixed position *x*, 1 ≤ *x* ≤ *k*, in *v* that are available for PS-node assignment. Also for a given visit to an rf-node *v* during TFS-alignment: let *R*(*V*′, *E*′) represent the PS-graph constructed so far; let 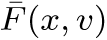 be those features that are already assigned to a PS-node and implicated for alignment in the current bundle; let 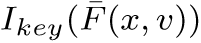 give the instance keys of those features that are already assigned; and finally let *V*′(*f*) give the PS-node to which a feature has been assigned. If at position *x* there is more than one feature, 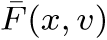, assigned to more than one PS-node, 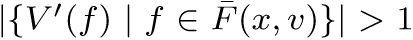, then PS-node assignments for each 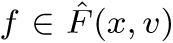 are determined by their respective instance key. In this case the instance keys of the unassigned features are compared to the instance keys of the assigned features.

Let *f* be an unassigned feature 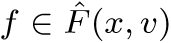 that we wish to assign, and *i* be its instance key 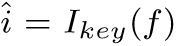; the assignment of *f* is determined by finding an instance key 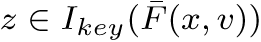 that has the largest intersection with i. If there is a tie between instance keys, then the one representing the PS-node with the most assignments is chosen. If when attempting the assignment of *f* a shift collision occurs, a new node in the PS-graph is created to which *f* is assigned. The r-component of the instance key helps to discriminate between available assignments when long stretches of repeat characters are present. Edges in the PS-graph are created between PS-nodes that have feature assignments which are consecutive in some *p* ∈ *P.* Through this procedure all featuress with the same instance key are guaranteed to be assigned to the same PS-node, and the full-path property of the PS-graph is maintained.

Shift collisions occur when bundling information attempts the alignment, in context *C*_*G*_ or *C*_*P*_, of two parts of the same sequence due to repeat characters. In the case of *C*_*G*_, when two bundles suggest the alignment of two sequences from the same genome, it can be indicative of a relative rearrangement that may be a fusion, fission, translocation, or misassembly. When a collision happens at the ends of two sequences from the same genome, it can be indicative of a missed join. In terms of the TFS-alignment procedure, when collisions occur between two different sequences in the same genome, no structural change is made to the graph. Panaconda creates a record of all such collisions, and the PS-nodes are marked as being involved in a collision. When collisions occur the type of event it implies will often depend on the “threading” pattern that results. In Figure 4, a graph of 42 *Brucella suis* is shown with a large translocation between replicons in two of the organisms. In this case, the two collision points anex a region that has moved from one chromosome to another.

**Fig. 4.**
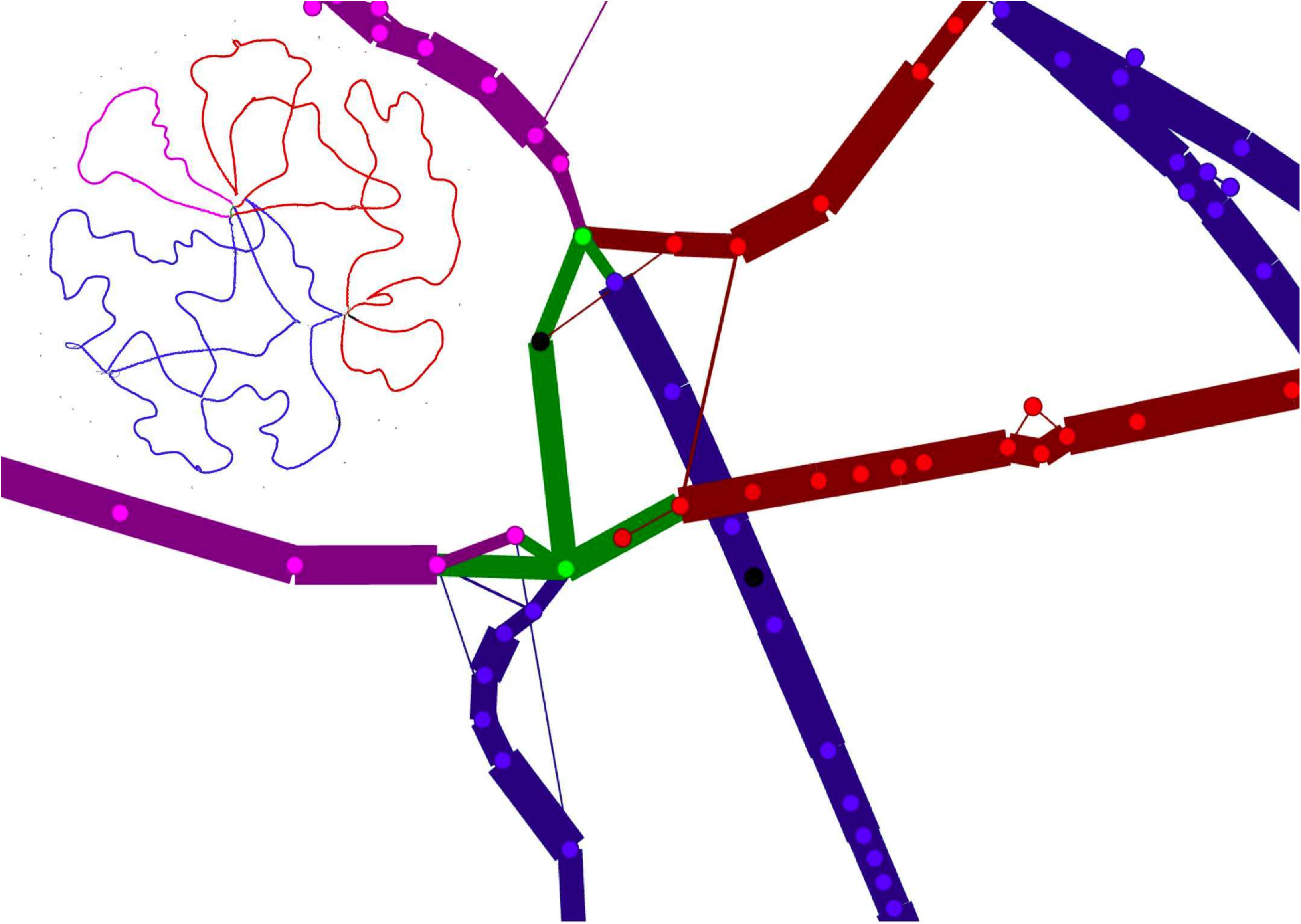
The region shown in fusia is typically found on chromsome 2 of Brucella suis. In the example shown a transloation has occurred in strains 1330 and VBI22 placing the region on chromosome 1. A zoomed out view is shown in the top left of the image.

In some cases a collision can occur that tries to align a sequence *to itself* which does not stem from repeat sequences. When this occurs, it is because there is a relative rearrangement in the alignment resulting from an inversion. In Figure 5 we show an inversion in a graph of 2 *Brucella melitensis* strains, 1 *Brucella microti,* and 1 *Brucella abortus.* The inversion is known in *Brucella abortus* bv. 1 str. 9-941 (Tsoktouridis *et al.,* 2003) and occurs on NC_003318, at between BMEII1009 hypothetical protein and BMEII0292 a putative Heme-regulated two-component response regulator.

**Fig. 5.**
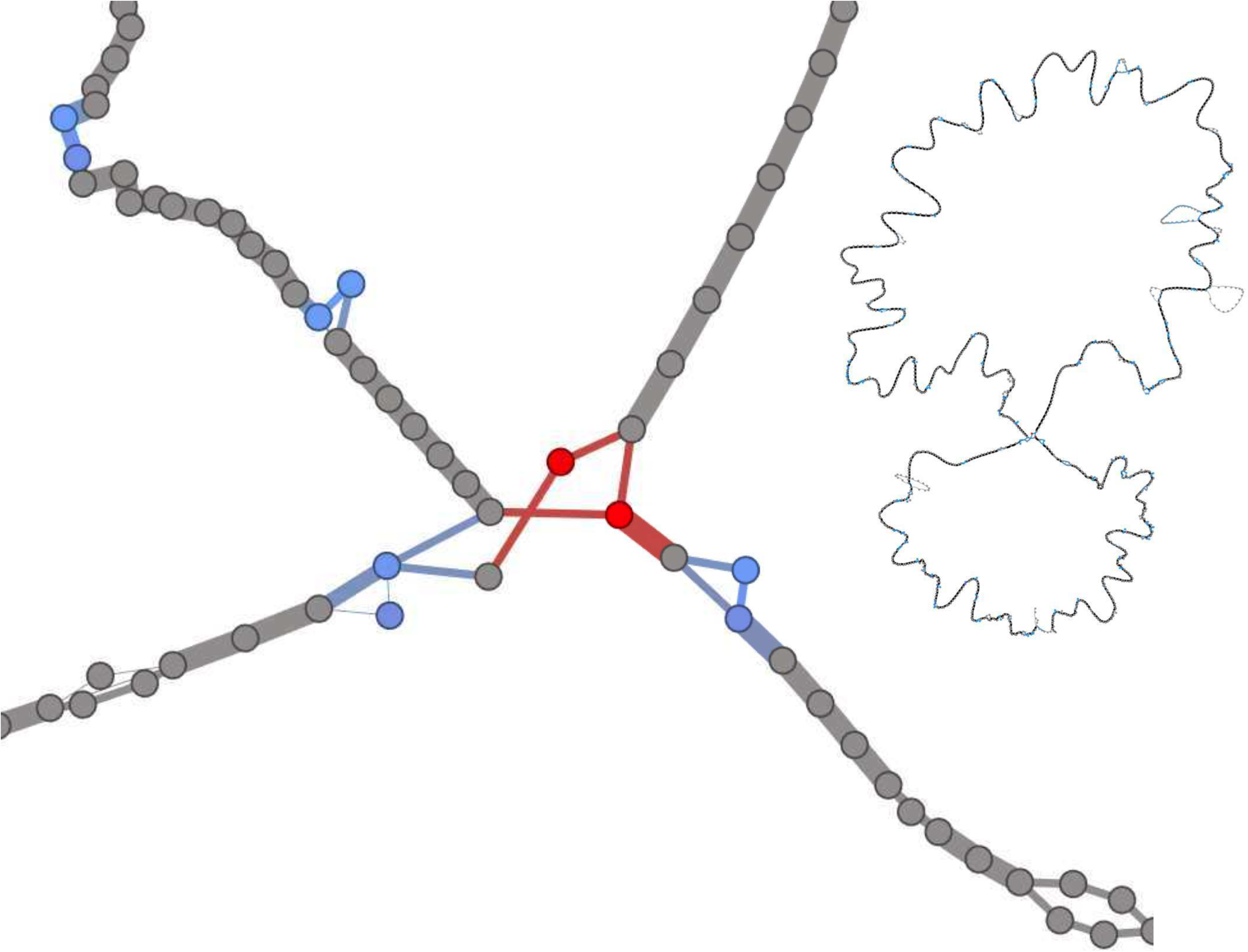
A zoomed (left) and high level perspective (right) on an inversion in Brucella abortus. The inversion collision is shown in red and shift collisions are shown in blue.

## 3 Results

Panaconda generates the pan-synteny graph, and optionally the RF-graph, in GEXF format. This makes the resulting alignment portable and usable in number of other programs, including Gephi (Bastian *et al*., 2009), NetworkX (Schult and Swart, 2008), and Cytoscape. Each node is labeled with the genome, sequence, and feature IDs assigned as part of the alignment process. Because the number of genomes represented in a graph (hundreds possibly thousands of genomes), the amount of metadata associated with the graph can be quite large. For pan-genome graphs in general, an index lookup scheme using a BGZF compressed backing store may improve portability. Panaconda’s graph creation component is implemented in Python using NetworkX. The component for laying out the graph is written in Java and is based on the Gephi toolkit. The visualization component is based on the Gexf-JS software, a browser based interactive GEXF viewer, and is written in Javascript and HTML.

### 3.1 Visualization

Multiple sequence alignment is a well researched and difficult problem (Darling *et al*., 2004; Chenna *et al*., 2003; Edgar, 2004), especially when dealing with diverse sequences that have complex evolutionary relationships (Minkin *et al*., 2013). Similarly, synteny has been explored often using line graphs (Arjona-Medina and Trelles, 2016; Alekseyev and Pevzner, 2009). In both types of analysis, the traditional grid based layout does not scale well with increasing numbers of sequences ((Lin *et al*., 2212). By using graphs, we are able to represent complex rearrangements, and hotspots of evolutionary change without sacrificing detail. The default Gexf-JS shows node attributes when clicking on a node and shows node labels when zoomed in. We have modified the base Gexf-JS viewer to expect and display a table of genome, sequence, and feature IDs. We have also added the ability of dynamic path highlighting for sequences and genomes.

As seen in Figure 6, when clicking in the node attribute table on a genome or sequence ID, all nodes and edges corresponding to that ID are highlighed. In this case, it allows us to see a suggested scaffolding for the unfinished genome *Mycobacterium tuberculosis* SP21 against *Mycobacterium tuberculosis* H37Rv. We have also configured Panaconda to calculate a diversity quotient based on the genus or species listed in the input annotaion. Based on the selected taxon level and the resulting diversity quotient, nodes are colored from yellow to red from low to high. The edge weight is set to be the fraction of genomes incident on that edge. With this difference in visual encoding, it is possible to see regions where certain species or genera are overrepresented. A command line option is provided to automatically lay out the pan-synteny graph that is generated. After trying a number of combinations, we have found that the multilevel agglomerative edge bundling (Gansner *et al*., 2011) implemented in older versions of Gephi works best. This is followed by Force Atlas 2 (Jacomy *et al*., 2014).

**Fig. 6.**
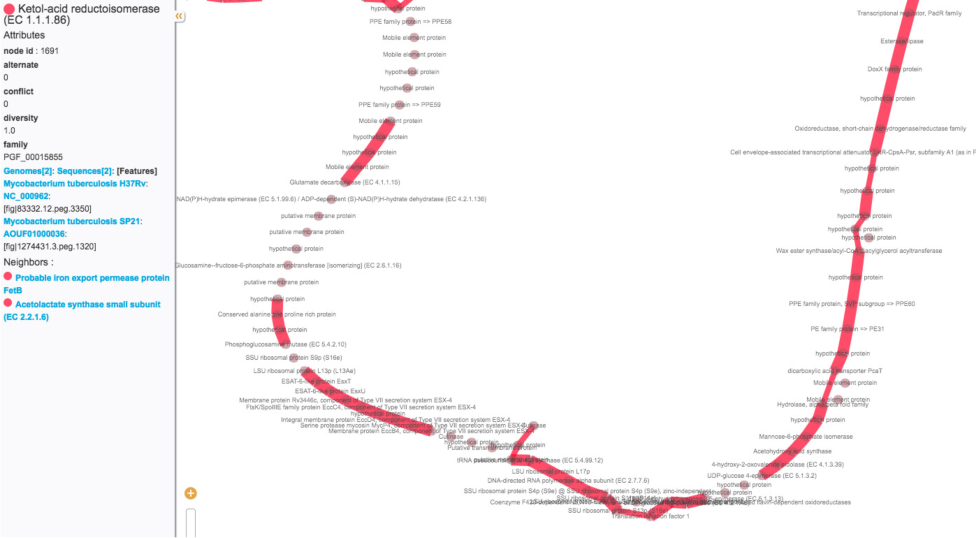
A potential scaffolding shown with two Tuberculosis strains H37Rv and SP21.

Evolutionary hotspots relative to the genomes in *G* can be recognized by a high degree of splitting of a strongly conserved block. Figure 7 shows several incidents of lateral transfer have occurred in the genus Brucella. Nineteen genes essential to the synthesis of lipopolysaccharide (LPS) and necessary to produce the smooth phenotype have been identified in the classically known Brucella (Godfroid *et al*., 2000; Gonzalez *et al*., 2008; Wattam *et al*., 2009). A study of atypical members of this genus (Wattam *et al*., 2012), and recent studies of Brucella isolated from amphibians (Soler-Llorens *et al*., 2016; Dahouk *et al*., 2017) have shown that phylogenetically more ancient members of the clade have different genes and build a different LPS. It has been shown that all Brucella share flanking regions and a tRNA gene where the inserts of these genes have occurred. The panaconda visualization built on the strains with the traditional Brucella O-antigen (B. abortus 2308, B. abortus 9-941, B. melitensis 16M, B. microti CCM 4915, B. inopinata BO1) and some strains isolated from amphibians (Brucella sp. 09RB8471, Brucella sp. 10RB9215 and Brucella sp. 09RB8910) show that there have been at least three separate insertions into this hotspot.

**Fig. 7.**
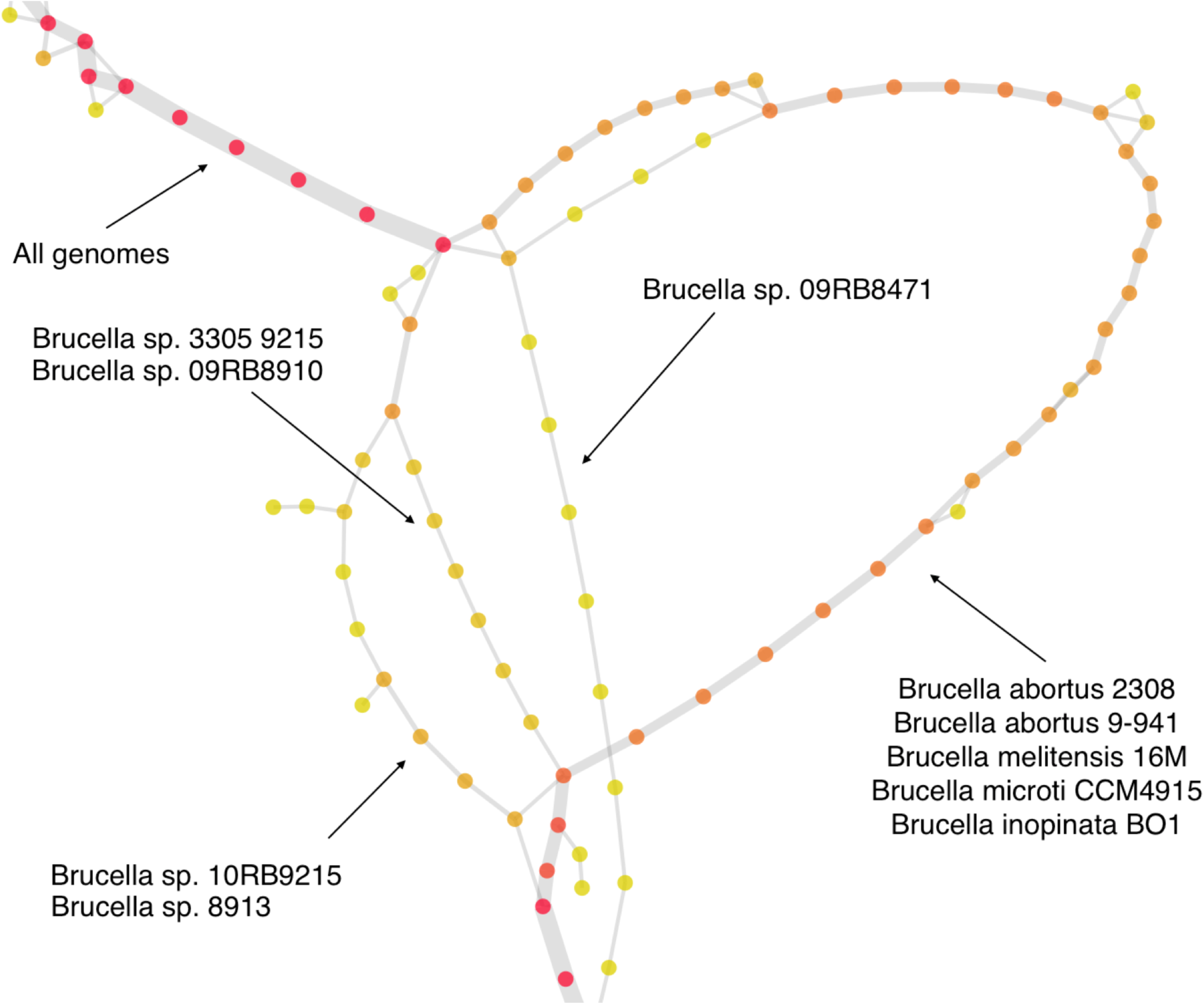
A highly divergent region in strains of Brucella

### 3.2 Performance

Because Panaconda takes advantage of previous computation invested in designating feature family assignments it runs fairly fast, taking under 3 minutes to align all available 565 sequences of Brucella from PATRIC (see Figure 8). Each visit to an rf-node in the TFS-alignment algorithm is guaranteed to align at least one feature *f* ∈ *F* giving an upper bound of 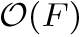 Intuitively, genomes that are more closely related are more likely to have more features co-incident on a given anchor node and thus have more features exhausted per visit than distantly related genomes. For example, aligning 100 Brucella genomes, a relatively well conserved genus, took 31 seconds and generated a PS-graph with 5,681 nodes compared to aligning 100 *Bacillus* genomes, a more divergent genus, which took 95 seconds and generated a graph with 116,289 nodes. All times were generated on a laptop with a 2.3 GHz Intel Core i7 processor and 16Gb of RAM.

**Fig. 8.**
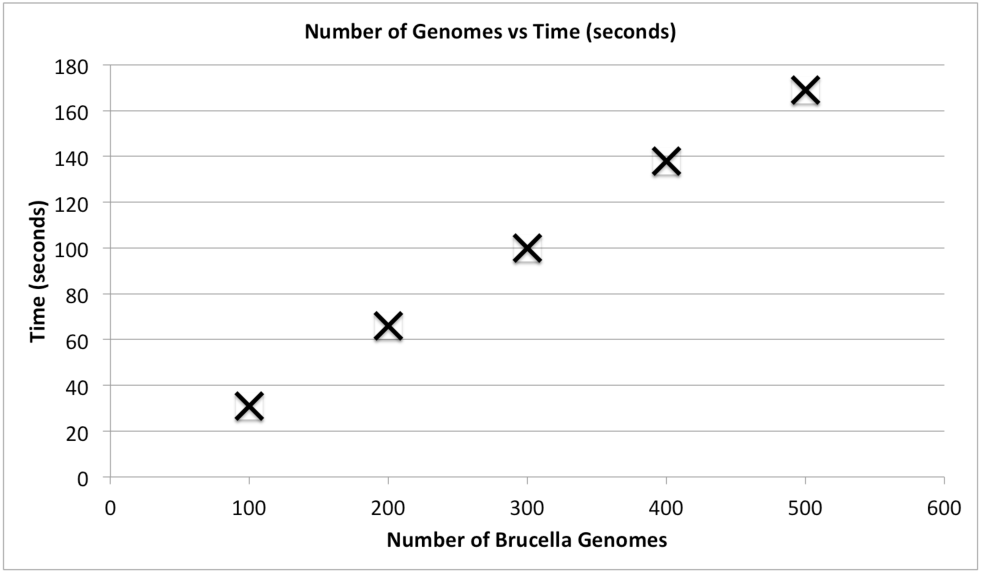
Run time of Panaconda in seconds on increasing numbers of Brucella genomes

## 4 Discussion

It is worth pointing out that pan-synteny graphs generated by Panaconda are closely related to the POA graphs introduced by Lee *et al*., 2002 in that they both model which positions in multiple sequences are aligned to one another; maintain the ordering of the positions within those sequences; fuses conserved letters into nodes as part of the alignment; and store sequence positions on the generated nodes. However, the approach of these two methods is somewhat different in that Panaconda creates a de Bruijn graph and uses it’s traversal to create the PS-graph from sequence conservation in any orientation, whereas POA requires that blocks of alignment occur in the same orientation. In their work on performing multiple sequence alignments on sequences with repeated and shuffled elements, Raphael *et al*., 2004 also use de Bruijn graphs to construct an alignment graph called an ABA graph. Besides the obvious difference in alphabet, pan-synteny graphs are different from ABA graphs in that the former parameterizes the the context of the alignment and does not contain cycles, with context setting *C*_*G*_ or *C*_*P*_, when following edge labels corresponding to the original sequence.

We find that Panaconda’s abstraction away from the traditional use of conservation of nucleotide sequence as the “glue” (Pevzner *et al*., 2004) that constitutes a homologous relationship can yield insightful results in prokaryotes. Panaconda algorithm uses multiple conserved k-mers of feature family annotations as the basis of its synteny recognition (multiplicity in the PS-graph greater than one). Like many synteny block finding algorithms, our approach requires a form of anchors. Unlike some previous methods (Peng *et al*., 2009), there is no requirement that a single anchor be present in all genomes. Unlike many synteny analysis algorithms (Pham and Pevzner, 2010; Minkin *et al*., 2013), the TFS-alignment method does not focus on alignment disruption due to loops or bulges (Peng *et al*., 2009). This can be attributed to the abundance of anchor k-mers owing to the large alphabet employed. As a result, no editing operations on sequences are needed, and the alignment is able to generate a graph where the full-path property is maintained.

We note that the default setting for Panaconda is a multiplicity count of one. In this default case, Panaconda creates and processes an RF-graph into a pg-graph which itself represents both syntenic blocks, in graph components with high multiplicity, and simple gene neighborhood relationships for those regions with only a single genome present. We assert that bacterial evolutionary relationships can be better understood when both synteny relationships and unique regions are analyzed together. Because of the complex rearrangements that can take place between and within bacterial genomes, explanatory methods that seek to characterize their relationships should be robust to large differences. We demonstrate that the pan-synteny graph is capable of locating and representing a wide range of evolutionary phenomena. By decoupling the comparison of genomes from strict nucleotide and amino acid sequence analysis, we are able to broaden or narrow the scope of the analysis depending on the unit abstraction chosen to create the alphabet. With our analysis, we supply examples that have three different types of alphabet provided by the PATRIC resource (Wattam *et al*., 2016): figfams (Meyer *et al*., 2009), pg-fams, and pl-fams (Davis *et al*., 2016).

## 5 Conclusion

This effort touches on several different research fronts: graph representation of genomes and their alignments, synteny block analysis, whole genome sequence alignment, pan-genome analysis, multiple sequence alignment, and genome rearrangement analysis. Though this approach was originally developed from a pan-genome perspective for prokaryotes, the methods involved have applicability to a wide range of topics. Elements of this work can be found in all topics described above, but we believe this approach represents a unique combination. Novel elements include the contextualization of synteny analysis both between and within multi-contig genomes. We also believe the algorithmic approach for discovering collision points has great value in the recognition of evolutionary relationships within a group of genomes.

Though synteny-block analysis has previously leveraged de Bruijn graphs for their determination, this work promotes the shift to a graphical model for the final representation of synteny relationships. Further we believe that such a shift combined with the abstract use of feature families not only adds utilityto pre-existing family databases, e.g. COGs and others, but can serve as a framework to speed up normally expensive operations such as phylogenetic tree construction and SNP analysis. This abstraction not only serves to make the resulting model approachable in terms of human cognition but also makes it's construction more resilient to distant and complex evolutionary relationships.

## Acknowledgements

The authors would like to acknowledge Eric Nordberg for his encouragement in pursuing this idea.

